# Mutational profiling of micro-dissected pre-malignant lesions from archived specimens

**DOI:** 10.1101/2020.04.05.026708

**Authors:** Daniela Nachmanson, Joseph Steward, Huazhen Yao, Adam Officer, Eliza Jeong, Thomas J. O’Keefe, Farnaz Hasteh, Kristen Jepsen, Gillian L. Hirst, Laura J. Esserman, Alexander D. Borowsky, Olivier Harismendy

**Author notes:** corresponding author: Olivier Harismendy.

## Abstract

**Background:** Systematic cancer screening has led to the increased detection of pre-malignant lesions (PMLs). The absence of reliable prognostic markers has led mostly to over treatment resulting in potentially unnecessary stress, or potentially insufficient treatment and avoidable progression. Importantly, most mutational profiling studies have relied on PML synchronous to invasive cancer, or performed in patients without outcome information, hence limiting their utility for biomarker discovery. The limitations in comprehensive mutational profiling of PMLs are in large part due to the significant technical and methodological challenges: most PML specimens are small, fixed in formalin and paraffin embedded (FFPE) and lack matching normal DNA.

**Methods:** Using test DNA from a highly degraded FFPE specimen, multiple targeted sequencing approaches were evaluated, varying DNA input amount (3-200 ng), library preparation strategy (BE: Blunt-End, SS: Single-Strand, AT: A-Tailing) and target size (whole exome vs cancer gene panel). Variants in high-input DNA from FFPE and mirrored frozen specimens were used for PML-specific variant calling training and testing, respectively. The resulting approach was applied to profile and compare multiple regions micro-dissected (mean area 5 mm^2^) from 3 breast ductal carcinoma in situ (DCIS).

**Results:** Using low-input FFPE DNA, BE and SS libraries resulted in 4.9 and 3.7 increase over AT libraries in the fraction of whole exome covered at 20x (BE:87%, SS:63%, AT:17%). Compared to high-confidence somatic mutations from frozen specimens, PML-specific variant filtering increased recall (BE:85%, SS:80%, AT:75%) and precision (BE:93%, SS:91%, AT:84%) to levels expected from sampling variation. Copy number alterations were consistent across all tested approaches and only impacted by the design of the capture probe-set. Applied to DNA extracted from 9 micro-dissected regions (8 PML, 1 normal epithelium), the approach achieved comparable performance, illustrated the data adequacy to identify candidate driver events *(GATA3 mutations, ERBB2* or *FGFR1 gains, TP53 loss)* and measure intra-lesion genetic heterogeneity.

**Conclusion:** Alternate experimental and analytical strategies increased the accuracy of DNA sequencing from archived micro-dissected PML regions, supporting the deeper molecular characterization of early cancer lesions and achieving a critical milestone in the development of biology-informed prognostic markers and precision chemo-prevention strategies.

## Background

For some cancer types, the wide-spread adoption of cancer screening has increased the detection of pre-malignant lesions (PML) [1]. Today, breast ductal carcinoma in situ (DCIS) comprises nearly ~25% of all breast cancer diagnoses in the United States [2]. In the case of DCIS, disease-specific guidelines recommend surgical excision and radiation, and endocrine risk reducing therapy. While treatment prevents rate of second events, it has not translated into increase survival rates, which are very high for most patients with DCIS, suggesting overtreatment of PML and highlighting a critical need to improve risk models and identify prognostic markers [1,3,4]. Current PML risk models rarely account for molecular biomarkers such as mutations or copy number alterations, which are seldom profiled. This is in large part due to technical challenges in profiling PML: specimens from PML biopsies are typically formalin fixed and paraffin embedded (FFPE) in their entirety, to verify absence of any invasive component. As a consequence, no fresh or frozen material is available for research. Moreover, many PMLs observed in absence of invasive lesions are very small or have low overall cellularity, sometimes less than a millimeter in diameter or containing fewer than 1000 cells. Hence, while FFPE specimens have successfully been used in high-throughput sequencing, the challenges posed by excessive formalin-induced deoxyribonucleic acid (DNA) damage – detailed below – are typically overcome by an increase in DNA input quantity [5–9], a solution not available for archival PML profiling. Thus, small FFPE PML specimens pose significant challenges in the generation of high throughput sequencing libraries and preclude the investigation of genetic biomarkers. To overcome these limitations, previous studies have been performed in fresh PML from areas adjacent to invasive disease instead of on pure PML, ignoring the vast majority of PML that are less likely to progress [10–15]. Profiling of pure PML in the absence of invasive disease is required to avoid such biases and thus necessitates methodology to work with archival FFPE specimens.

One of the main challenges of library preparation from damaged, low input DNA samples is to preserve the library complexity: the faithful and unbiased representation of all fragments in the starting DNA sample. Indeed the multiple steps of the library preparation, including the repair of the input DNA, the ligation of adapter, target enrichment and the multiple rounds of purification and PCR amplification can all act as bottlenecks, and introduce strong skews that will reduce the library complexity and eventually impact precision and recall of variant calling. Moreover, formalin is known to create adducts in the DNA and lead to spurious substitutions, which can be difficult to distinguish from true somatic variants, especially at low allelic fractions [16,17]. Finally, the most insightful prognostic biomarker studies of PML progression require long follow up to rely on actual outcome (recurrence, second events, survival) rather than proxy risk markers (grade, subtype, histological markers). As a consequence, most studies are retrospective and rely on old archived material without matching germline DNA sample, rendering the identification of high confidence somatic mutations more difficult. Hence, both technical and experimental challenges are hampering progress in PML mutational profiling.

Here we present the development of DNA library preparation and variant calling strategies specifically optimized for low abundance, damaged DNA, commonly extracted from PMLs. Using highly damaged DNA, we compared the effect of the input amount, the size of the captured genomic region, and library preparation strategy on the quality of coverage depth and breadth. We determined that library preparation using blunt-end (BE) adapter ligation strategy maximizes the library complexity down to 3 nanograms (ng) of input DNA and is compatible with whole exome capture. Using a set of DNA variants called from a frozen mirrored tissue specimen, we optimized the variant calling strategy to maximize its accuracy. We further demonstrated its validity on 10 DNA samples extracted from laser capture micro-dissected regions of PML or adjacent normal from DCIS of 3 independent patient FFPE specimen. We illustrate the utility of the approach to identify somatic mutations in candidate genes and characterize PML clonal heterogeneity within a specimen.

## Methods

### Sample collection and preparation

#### Test specimen

Mirrored frozen – FFPE tissue specimen from a HER2 positive invasive breast cancer was obtained from Asterand Biosciences (Detroit, MI). The length distribution of the DNA fragments was measured by capillary electrophoresis (Agilent BioAnalyzer) and used to calculate the DNA integrity number (DIN) between 1 (very degraded) to 10 (intact genomic DNA). *<u>DCIS specimen</u>:* FFPE blocks were obtained from UCSD Health Anatomic Pathology after surgical biopsy, excision or mastectomy. The UCSD institutional review board approved the retrospective study and waived the requirement for consent. Consecutive sections of the blocks were used for Hematoxylin-Eosin staining (N=1; 4 μM glass slide) then for Laser Capture Microdissection (LCM; N=3; 7 μM glass slide coated with polyethylene naphthalate – ThermoFisher #LCM0522). The slides were stored at −20°C in an airtight container with desiccant until ready for dissection (1 day to 3 months). The LCM sections were thawed and stained with eosin, sections were kept in xylene and dissected within 2 hours of staining. Laser Capture Microdissection was performed using the Arcturus Laser Capture Microdissection System. Matching regions from 3 adjacent sections were collected on one Capsure Macro Cap (Thermofisher), region size permitting. Post-dissection, all caps were covered and stored at – 20°C. *<u>DNA extraction and QC</u>:* The DNA was extracted from FFPE tissue using the QIAamp DNA FFPE tissue kit and QIAamp DNA Micro Kit (Qiagen) for the test specimen or LCM specimen, respectively. For the LCM sample, the membrane and adhering tissue were peeled off the caps using a razor blade and the peeled membrane was incubated in proteinase K digestion reaction overnight for 16 h at 56°C to maximize DNA yield after cell lysis and the elution was done in 20 μL. The extracted DNA was quantified by fluorometry (HS dsDNA kit Qbit – Thermofisher). All samples used in the study are described in Additional file 1: Table S1.

### Targeted sequencing

#### DNA fragmentation

DNA was sheared down to 200 base pairs (bp) using Adaptive Focused Acoustics on the Covaris E220 (Covaris Inc) following manufacturer recommendations with the following modifications: 50 μL of Low TE buffer in microTUBE-130 tubes (AT libraries) or 10 μL Low EDTA TE buffer supplemented with 5 μL of truSHEAR buffer using a microTUBE-15 (SS and BE libraries).

#### Library preparation

AT libraries were prepared with the SureSelect XT HS protocol (Agilent Technologies) extending the adapter ligation time to 45 minutes (min). After ligation, excess adapters were removed using a 0.8x SPRI bead clean up with Agencourt AMPure XP beads (Beckman Coulter), then eluted into 21 μL of nuclease-free water. SS libraries were prepared using the Accel-NGS 1S Plus DNA Library Kit (Swift Biosciences). Prior to the single-strand ligation protocol, 15 μL of fragmented DNA was denatured at 95°C for 2 min, then set on ice. The adaptase and extension steps were performed by kit specifications followed by a purification step using 1.2x AMPure XP, eluted into 20 μL of nuclease free water. The subsequent ligation step incorporates SWIFT-1S P5 and SWIFT-1S P7 adapters, followed by a 1x AMPure XP bead clean-up and elution into 20 μL of nuclease free water. BE libraries were prepared using the Accel-NGS 2S PCR-Free DNA Library Kit (Swift Biosciences). Repair I was followed by a 1x AMPure bead cleanup, Repair II was followed by a 1x PEG NaCl cleanup, and Ligation I (P7 index adapter) and Ligation II (P5 UMI adapter) were followed by a 0.85x PEG NaCl cleanups. Only Ligation II cleanup was eluted into 20 μL Low EDTA TE, the other cleanups proceeded directly into the next reaction. Adapters used in the study are summarized in Additional file 1: Table S2.

#### Pre-Capture PCR amplification

Ligated and purified libraries were amplified using KAPA HiFi HotStart Real-time PCR 2X Master Mix (KAPA Biosystems). AT libraries were amplified with 2 μL of KAPA P5 primer and 2 μL of SureSelect P7 Index primer. SS libraries were amplified with 5 μL of SWIFT-1S P5 Index and P7 Index primers. BE samples were amplified with 5 μL of KAPA P5 and KAPA P7 primers. The reactions were denatured for 45 seconds (sec) at 98°C and amplified 13-15 cycles for 15 sec at 98°C, for 30 sec at 65°C, and for 30 sec at 72°C, followed by final extension for 1 min at 72°C. Samples were amplified until they reached Fluorescent Standard 3, cycles being dependent on input DNA quantity and quality. PCR reactions were then purified using 1x AMPure XP bead clean-up and eluted into 20 μL of nuclease-free water. The resulting libraries were analyzed using the Agilent 4200 Tapestation (D1000 ScreenTape) and quantified by fluorescence (Qubit dsDNA HS assay). Primers used in the study are summarized in Additional file 1: Table S2.

#### Targeted Capture Hybridization and Post-Capture PCR

Samples were paired and combined (12 μL total) to yield a capture “pond” of at least 350 ng, and supplemented with 5 μL of SureSelect XTHS and XT Low Input Blocker Mix. The baits for target enrichment consisted of either Agilent SureSelect Clinical Research Exome panel (S06588914), Human All Exon V7 panel (S31285117) or Cancer All-In-One Solid Tumor (A3131601). The hybridization and capture was performed using Agilent SureSelect XT HS Target Enrichment Kit following manufacturer’s recommendations. Post-capture amplification was performed on the beads in a 25 μL reaction: 12.5 μL of nuclease-free water, 10 μL 5x Herculase II Reaction Buffer, 1 μL Herculase II Fusion DNA Polymerase, 0.5 μL 100 millimolar (mM) dNTP Mix and 1 μL SureSelect Post-Capture Primer Mix. The reaction was denatured for 30 sec at 98°C, then amplified for 12 cycles of 98°C for 30 sec, 60°C for 30 sec and 72°C for 1 min, followed by an extension at 72°C for 5 minutes and a final hold at 4°C. Libraries were purified with a 1x AMPure XP bead clean up and eluted into 20 μL nuclease free water in preparation for sequencing. The resulting libraries were analyzed using the Agilent 4200 Tapestation (D1000 ScreenTape) and quantified by fluorescence (Qbit – ThermoFisher)

#### RNA-sequencing

Expression profiling was performed on select dissected PML regions: 1A, 2A2, 2B and 3C. RNA library preparation was performed with SMART-3Seq, a 3’ tagging strategy specifically designed for degraded RNA directly from FFPE LCM specimen [18]. Read count data was obtained using a dedicated analysis workflow https://github.com/danielanach/SMART-3SEQ-smk. Count data was then normalized for read depth and scaled by a million to give transcripts per million (TPM) counts.

#### Sequencing

All libraries were sequenced using on the HiSeq 4000 sequencer (Illumina) for 100 cycles in Paired-End mode. Libraries with distinct indexes were pooled in equimolar amounts. The sequencing and capture pools were later deconvoluted using program bcl2fastq [19].

### Data analysis

#### Sequencing reads processing and coverage quality control

Sequencing data was analyzed using bcbio-nextgen (v1.1.6) as a workflow manager [20]. Samples prepared with identical targeted panels were down-sampled to have equal number of reads using seqtk sample (v1.3) [21]. Adapter sequences were trimmed using Atropos (v1.1.22), the trimmed reads were subsequently aligned with bwa-mem (v0.7.17) to reference genome hg19, then PCR duplicates were removed using biobambam2 (v2.0.87) [22–24]. Additional BAM file manipulation and collection of QC metrics was performed with picard (v2.20.4) and samtools (v1.9) [24]. The summary statistics of the sequencing and coverage results are presented in Additional file 1: Table S3.

#### Copy number analysis

Copy number alterations (CNAs) were called using CNVkit (v0.9.6) [25] using equal sized bins of ~250 bp. Any bins with log_2_ copy ratio lower than −15, were considered artifacts and removed. Breakpoints between copy number segment were determined using the circular binary segmentation algorithm (p < 10^-4^)[26]. Low quality segments were removed from downstream analysis (less than 10 probes, biweight midvariance more than 2 or log_2_ copy ratio confidence interval contains 0). Copy number genomic burden was computed as the sum of sizes of segments in a gain (log_2_(ratio)>0.3) or loss (log_2_(ratio)<-0.3) over the sum of the sizes of all segments. The summary statistics of CNA calling on the test specimen are reported in Additional file 1: Table S4. Chromosomal arm gains and losses were called when more than half their total length was involved in a gained and lost segment, respectively. Gene copy number estimates were assigned based on the segment that covered the gene. For the *<u>test specimen</u>,* if more than one segment covered a gene then the higher confidence segment was used. For the *<u>DCIS specimen</u>* copy number alterations were determined for all autosomal genes containing at least 3 bins whose segments were covered by at least 110 probes (N=17,750 total). Additionally, due to the imprecision of segmentation breakpoints, any genes with a breakpoint identified in one region of a patient, were removed from the comparison to other regions. Loss of heterozygosity (LOH) was called for segments with B-allele frequencies lower than 0.3 or greater than 0.7.

#### Variant calling and initial filtering

Single nucleotide variants (SNVs) and short insertions and deletions (indels) were called with VarDictJava (v1.6.0), and Mutect2 (v2.2) [27,28]. Variants were required to fall within a 10 bp boundary of targeted regions that overlapped with RefSeq genes (v 109.20190905). A publicly available list of variants observed in a pool of normal DNA exome sequencing was obtained from the GATK resource (https://console.cloud.google.com/storage/browser/details/gatk-best-practices/somatic-b37/Mutect2-exome-panel.vcf) and used to eliminate artifacts and common germline variants. Only variants called by both algorithms were considered (ensemble calling). These ensemble variants were then subjected to an initial filtering step with default bcbio-nextgen tumor-only variant calling filters listed in Additional file 1: Table S5. Functional effects were predicted using SnpEff (v4.3.1) [29]. The resulting variants are referred to as <u>raw ensemble variants</u>.

#### Germline variant filtering

In absence of a matched normal control for both test (frozen and FFPE) and DCIS specimens, somatic mutations were prioritized computationally using the approach from the bcbio-nextgen tumor-only configuration then additionally subjected to more stringent filtering [30]. Briefly, common variants (MAF>10^-3^) present in population databases – 1000 genomes (v2.8), ExAC (v0.3), or gnomAD exome (v2.1) – were removed unless in a tier 1 gene from the cancer gene consensus and present in either COSMIC (v68) or clinvar (20190513) [20,31–35]. Additionally, variants were removed as likely germline if found at a variant allelic fraction (VAF) greater or equal to 0.9 in non-LOH genomic segments – as determined by CNA analysis (above). The remaining variants are referred to as <u>candidate somatic mutations</u>.

#### Analysis specific filtering of candidate somatic mutations

Additional filtering was implemented in a context specific to the analysis presented. *1) <u>Calling “gold standard” mutations from the frozen test specimen</u>:* the candidate somatic mutations in the DNA of the frozen test specimens were filtered for high-quality variants: ensemble quality score greater than 175, average number of read mismatches less than 2.5, position covered by at least 25 reads, mean position in read greater than 20, microsatellite length less than 5, and VAF more than 0.14. This resulted in 247 SNVs and 10 indels. 2) *<u>Benchmarking mutations in FFPE test specimens</u>:* mutations in the DNA of the FFPE test specimens required specific filtering due to the abundant low-frequency damage as well as lower coverage depth. The following parameters were used: position covered by at least 5 reads, mapping quality more than 45, mean position in read greater than 15, number of average read mis-matches less than 2.5, microsatellite length less than 5, tumor log odds threshold more than 10, Fisher strand bias Phred-scaled probability less than 10 and VAF more than 0.14. The accuracy of resulting DNA variants from the test FFPE specimen was measured against the set of “gold standard” variants from the mirrored frozen specimen using vcfeval by RTG-tools (v3.10.1), using variant ensemble quality as the score [36]. The results of the benchmarking analysis are reported in Additional file 1: Table S6. 3) *<u>Profiling dissected regions from the DCIS specimen</u>:* in addition to the filtering of FFPE candidate somatic mutations presented above, the following steps were implemented. Any variants found at high VAF (>0.9) in non-LOH segments in one region were also excluded from the variants from all other regions of same patient. Candidate somatic mutations with ensemble quality score lower than 115 were excluded, corresponding to the optimal F-score obtained for low-input BE libraries in the benchmarking analysis. To the exception of few well-described hotspot mutations in breast cancer *(PIK3CA, TP53, GATA3),* somatic mutations identified in more than one patient were removed.

#### Clonality analysis

To allow the analysis of clonal relationships between regions of the same patient, the coverage depth of each allele at any remaining mutated position in any region was extracted using Mutect2 joint variant caller on the sets of aligned reads from each region. In order to call a mutation either absent or present in a region, we used a Bayesian inference model specifically designed for multi-region variant calling [37]. Treeomics (v1.7.10) was run with the default parameters except for e=0.02.

## Results

### Evaluation of targeted sequencing approaches for low input FFPE DNA

Regions of PMLs on a histological section can contain as few as 500 cells, corresponding to 3.3 ng of haploid DNA. The expected reduced DNA extraction yield can be mitigated by combining regions matched across sequential sections. Hence, optimizing targeted sequencing down to 3 ng of input DNA is a reasonable objective to identify high confidence mutations in PML. To develop the methodology, a test DNA sample was extracted from a 4 year-old FFPE HER2 positive breast invasive carcinoma which showed significant fragmentation (DNA integrity number of 2.4), likely representative of DNA extracted from old archived specimens. High-throughput sequencing libraries were prepared using traditional A-tailing adapter ligation protocol optimized for low-input, damaged DNA (referred to as AT method – Figure 1A) with decreasing amount of input DNA from 200 down to 10 ng. After capture of the whole-exome by hybrid selection, the DNA libraries were amplified and sequenced. The performance was evaluated in comparison to whole exome data generated from 200 ng of DNA extracted from a mirrored frozen tissue specimen [38]. Libraries generated from 200 or 50 ng FFPE DNA achieved reasonable coverage, with nearly all targeted bases covered at least 20-fold (referred to as Cov20). In contrast, the sequencing libraries from 10 ng FFPE DNA lacked complexity (79% read PCR duplicates), resulting in 17% Cov20 after elimination of the duplicate reads (Figure 1B-C). Such poor performance at 10 ng, precluded us from further decreasing the DNA input and suggested that perhaps a smaller capture panel, restricted to cancer genes (710 kb total size) would elicit the goal of 3 ng input. Unfortunately, 3 ng AT libraries sequencing with a cancer panel had a high percentage of duplicate reads (83%) and 1.6% Cov20 (Figure 1D-E). Hence, the consistent poor performance of both low input strategies (3 ng and 10 ng) irrespective of the capture panel size suggests that the bottleneck reducing the library complexity originates upstream of the targeted capture, which led us to examine the initial ligation of the sequencing adapters.

**Figure 1.**
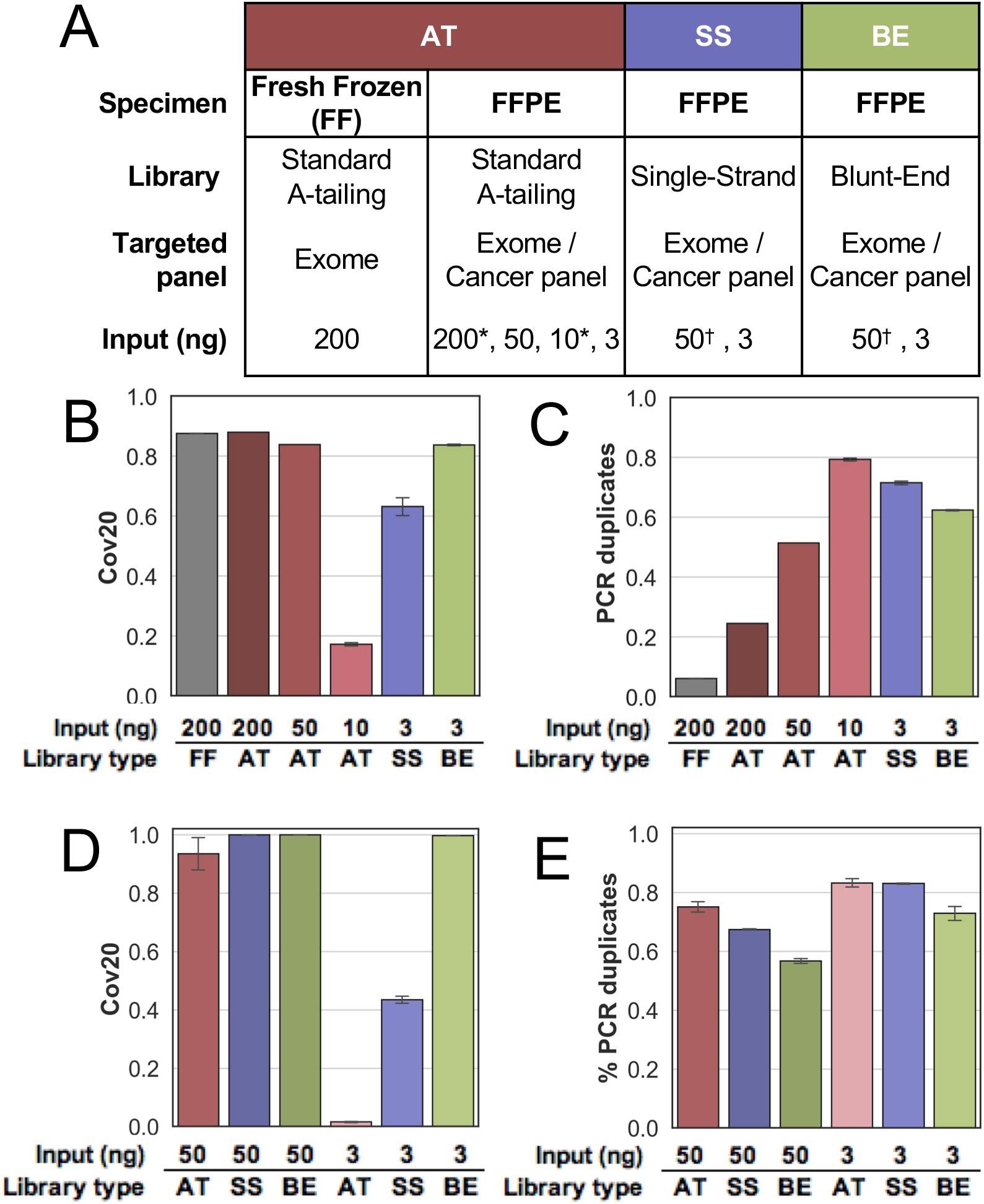
Benchmarking results for sequencing performance. **(A)** Experimental design for performance evaluation using a test DNA specimen (* Exome and † Cancer panel). **(B-C)** Fraction of targeted bases covered by a minimum of 20 reads (B) and fraction of PCR duplicates (C) observed in whole exome sequencing. **(D-E)** Fraction of targeted bases covered by a minimum of 20 reads (D) and fraction of PCR duplicates (E) observed in cancer panel sequencing. All error bars represent standard deviation.

Library preparation methods which utilize alternate ligation techniques have previously been shown to increase the complexity of the library notably for the analysis of cell free DNA [39,40], ancient DNA [41–43] and other applications [44,45]. We therefore evaluated blunt-end ligation (BE) and single strand ligation (SS) library preparation strategies to increase the number of input DNA fragments incorporated in the library and enhance the resulting complexity and usable sequence coverage (Figure 1A, Additional file 2: Figure S1). The cancer panel capture of BE libraries prepared from 3 ng of test DNA showed a reduced percentage of duplicate reads compared to AT libraries with 10 ng input (73% vs 83% respectively Figure 1E), leading to a dramatic increase in Cov20 (99.7% vs 2%) and bringing it to levels comparable to high input (50 ng) AT libraries (Figure 1D), albeit with higher duplicate rates. In turn, the SS library offered a lesser, but measurable, improvement over AT library for low input (Figure 1D-E). The superiority of the BE and SS strategies were further confirmed using whole exome capture with BE and SS libraries resulting in 84% and 63% Cov20 at low input (3 ng), respectively, higher than 17% observed for low input (10 ng) AT libraries (Figure 1B). Compared to the cancer panel capture, the improvements of these alternative ligation strategies on whole exome sequencing were milder but remained remarkable and, in the case of BE strategy, likely to support more sensitive mutational profiling of archived PMLs.

### Somatic mutation profiling from low input FFPE DNA

At a minimum, both SNVs and indels are necessary to evaluate the mutational landscape of PML. Unfiltered SNVs identified in FFPE DNA showed both a high overall abundance (422,322) of low variant allelic fraction substitutions (VAF <5%) and bias of C to T substitutions (53%), which is expected from the cytosine deamination resulting from formalin fixation [16,17]. In contrast, low VAF variants from frozen DNA were much lower in abundance (175,364) but contained a high-prevalence (52%) of C to A substitutions consistent with previously reported 8-oxoguanine damage observed in frozen samples (Additional file 2: Figure S2) [46]. We hypothesized that we could discriminate against artifactual FFPE variants in the test specimens using stringent filtering criteria including high strand bias, low allelic fraction and poor concordance between multiple variant callers in order to call accurate somatic variants in FFPE preserved PML (Methods).

First, we established a set of benchmarking somatic mutations from the test DNA extracted from a mirroring frozen tissue specimen [38]. Its whole exome sequence resulted in 247 SNVs and 10 indels that were used to measure performance of the variant calling from the FFPE libraries generated above. Prior to filtering, the analysis of variants from low input AT libraries resulted in an average of 7,475 false positive somatic mutations. In contrast, the analysis of variants from BE and SS libraries resulted in an average of 1,967 and 3,137 false positive mutations, respectively (Figure 2A). We developed additional filtering criteria, trained on variants from the high-input (200 ng) FFPE library and used variants called from a publicly available panel of normal samples to remove additional artifacts. This approach considerably reduced the fraction of false positives (Figure 2B), increasing precision from less than 20% to 84%, 93% and 91% for the AT, BE and SS low input libraries, respectively (Figure 2C-D). The variant recall increased from 75% in the AT low input library to 85% and 80% for the BE and SS low-input libraries, respectively, consistent with differences observed in Cov20 (Additional file 1: Table S6). Importantly, these values were similar to the theoretical maximum values obtained from downsampling the frozen sample itself to an equivalent number of reads (90% precision, 88% recall), indicating that the differences mainly comes from sampling rather than from technical artifacts. The improvement in accuracy was similar for the small number of indels (Additional file 1: Table S6). These data suggest that FFPE specific filtering of ensemble variant calls paired with BE library preparation enables accurate clonal somatic SNV and indel variant calling of whole exome sequencing data from 3 ng FFPE test DNA.

**Figure 2.**
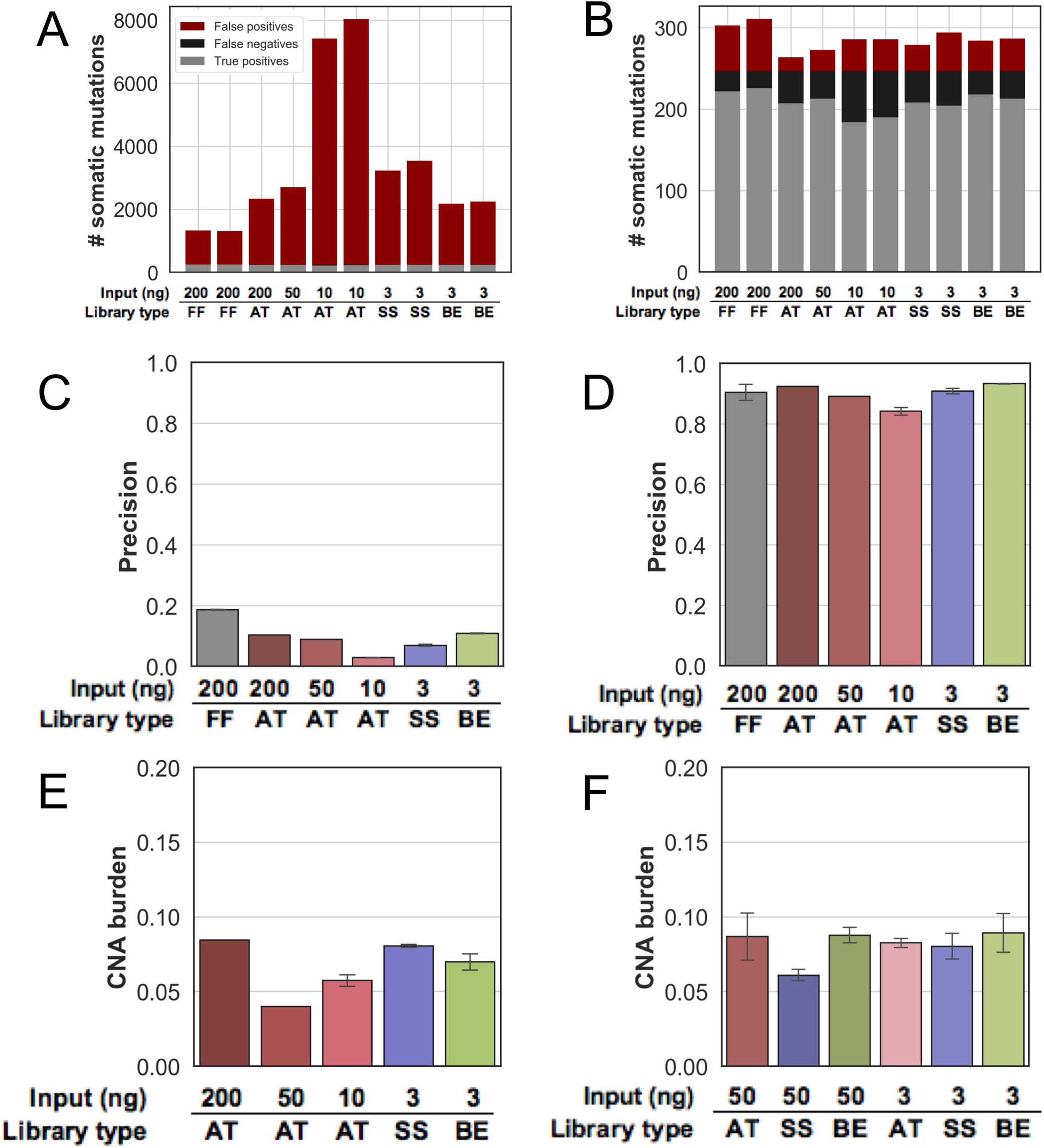
Benchmarking results for variant calling. **(A-B)** Count of total variants from whole exome sequencing, separated by false positives (red), false negatives (black) and true positives (grey), before (A) and after (B) PML specific filtering. (**C-D**) Exome variant calling precision for various library preparation strategies and amount of input DNA (x-axis) before (C) and after (D) PML specific filtering. (**E-F**) Fraction of the genome involved in a copy number alteration (CNA burden – y axis) for all exome (E) and cancer panel (F) library preparation strategies and DNA input amounts. All error bars represent standard deviation.

In contrast to SNVs and indels, Copy Number Alterations (CNA) can be accurately identified using lower coverage depth whole genome sequencing data, though has been more challenging in targeted sequencing [47,48]. We evaluated the ability of all input quantity and library preparation methods to reliably identify CNA (Methods). The genome-wide CNA burden obtained from low input SS and BE library was consistent with the one of high-input (200 ng) AT libraries (8.1%, 7% and 8.4% respectively – Figure 2E and Additional file 1: Table S4), and consistent with the results of the cancer panel sequencing and the lower CNA burden observed in the 10 and 50 ng AT libraries are still within the expected confidence interval (Additional file 2: Figure S3). The resulting level of copy number gains and losses estimated for each chromosome arms and more focal areas were highly reproducible between all tested library preparation strategies (Additional file 2: Figure S4-S5). Additionally, the copy number status of known cancer genes was also consistent between exome and cancer panel, including the expected *ERBB2* amplification which was correctly determined in all cases. In a few instances, the denser tiling of the exome probes helped identify a copy number breakpoint missed in the cancer panel, resulting in discordant copy number estimate for a few genes (Additional file 2: Figure S6). This suggests that the input amount and quality of the sample has little impact on the accuracy of the copy number profiling, albeit this observation was limited to a specimen with few CNAs.

### Mutational profiling of breast PML

To validate the optimized targeted sequencing and somatic variant calling on PMLs, we collected archived tissue specimens from 3 patients diagnosed with breast ductal carcinoma in situ (DCIS – 1 low grade, 2 high grade) without evidence of invasive disease. A total of 8 PML regions and one normal breast epithelium region (patient 3, region 3N) were isolated by laser capture microdissection (LCM – Figure 3A). The dissected regions had a mean area of 5 mm^2^ and, combined over three adjacent sections, contained an average of 80,000 cells (Figure 3B). For one large DCIS region, adjacent sections (patient 2, regions 2A1 and 2A2) were processed independently for replication. For each region, between 1.4 and 21 ng of DNA were extracted and used to prepare exome libraries using the BE method. The rate of duplicate reads was between 32 and 82%, and, as expected, inversely correlated with the amount of input DNA (Additional file 2: Figure S7). The resulting mean coverage depth was comprised between ~2 and 45 fold, which is sufficient for accurate detection of CNA, but likely limiting the sensitivity to identify mutations, particularly in patient 2.

**Figure 3.**
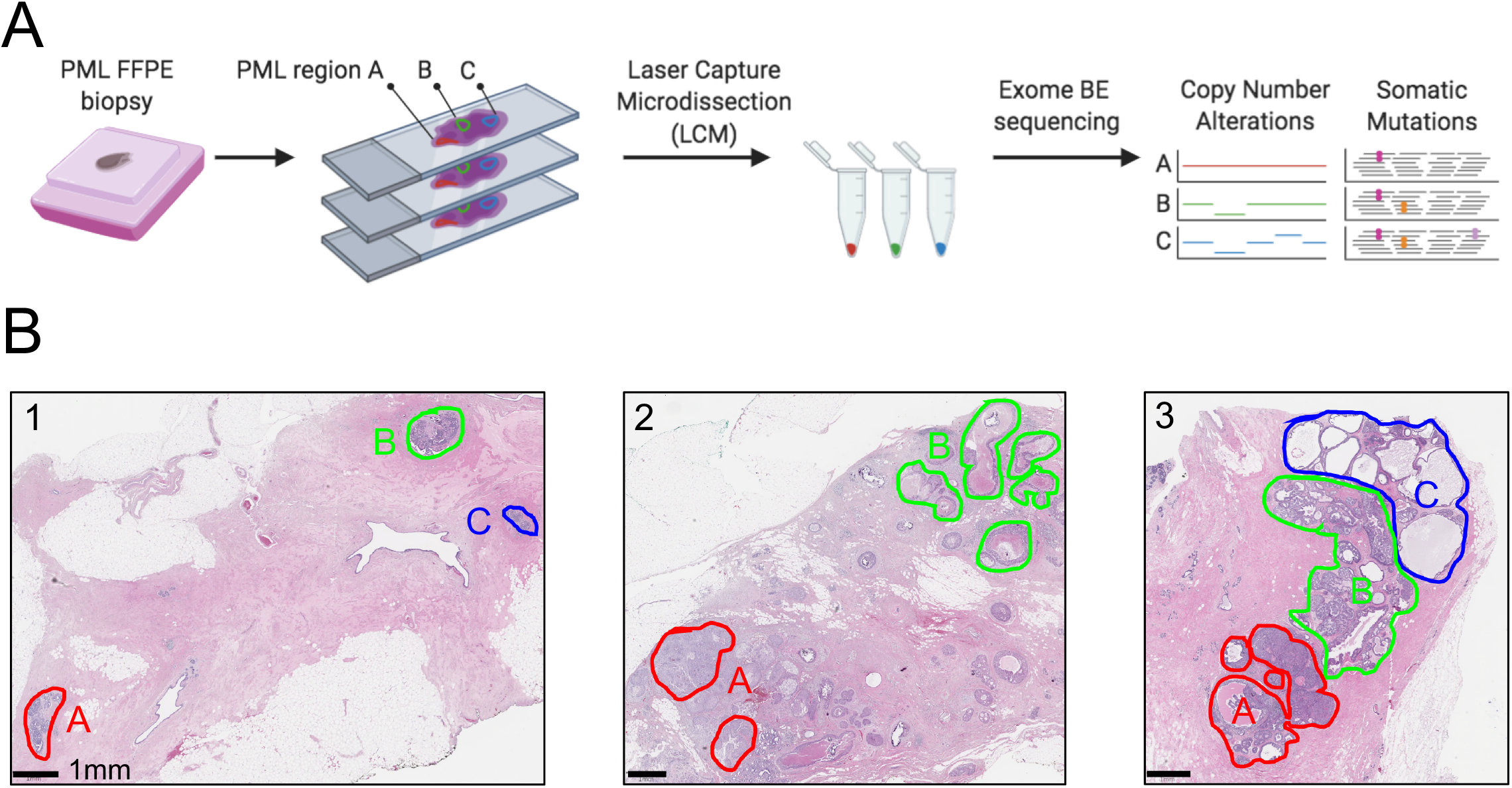
Overview of the PML regional sequencing strategy. **(A)** Overall experimental and analytical workflow of the validation study. Images created with BioRender.com and printed with permission. **(B)** Images showing the Hematoxylin and Eosin stained sections from the three DCIS patient studied. Dissected PML regions are highlighted in color to the exception of region 3N consisting of multiple areas of normal epithelium outside the selected field of view.

Between 7% and 43% of the regions’ genomes were involved in CNAs, predominantly through copy number losses (Figure 4A). With the exception of region 2B, the CNA burden was consistent between regions of the same patient and minimal in normal breast epithelium (<5%). A total of 18 chromosome arms were affected by copy number changes in at least one DCIS region. None of these were altered in the normal breast epithelium (Figure 4B). Some of the chromosome arm losses identified, such as 6q, 8p, 16q, 17p or 22q are frequent in DCIS [49]. Within patient 1 and 3, all regions have consistent CNA suggesting a common clonal ancestor. In contrast, regions 2A and 2B have different number of arms altered (5 vs 13, respectively) with only 3 in common. Both regions featured high nuclear grade, but only 2B showed comedonecrosis, a marker of more advanced and worse prognosis DCIS. Region 2B was the only region affected by chromosome arm gains at 8q, 20q and 21q. The absence of these gains in 2A as well as its additional losses of 4p and 9p, suggest the independent clonal evolution of 2A and 2B. Finally, all PML regions from patient 2 and 3 displayed a loss of 17p, generally associated with *TP53* loss of heterozygosity (LOH) frequently observed in high-grade specimens. At a higher resolution, out of 98 cancer genes evaluated, 38 had a CNA in at least one region (Figure 4C). The most notable ones were the amplification of *ERBB2* in all regions of patient 2, amplification of *FGFR1* in all regions of patient 3 and loss of *TP53* in patient 2 and 3. This latter observation was consistent with 17p LOH and confirmed by change in B-allele frequency at heterozygous SNPs (Additional file 2: Figure S8). The relative gene expression levels measured for 3 genes in 4 matching regions were consistent with their copy number in 10/12 cases, with possible discrepancies due to spatial variation in histology or transcriptional regulation (Additional file 2: Figure S9) [18]. Similar to chromosome arms, none of the genes were altered in the normal epithelium and most had consistent copy numbers between regions of the same patient, with the exception of 12 genes distinguishing regions 2A and 2B and further supporting separate clonal evolution.

**Figure 4.**
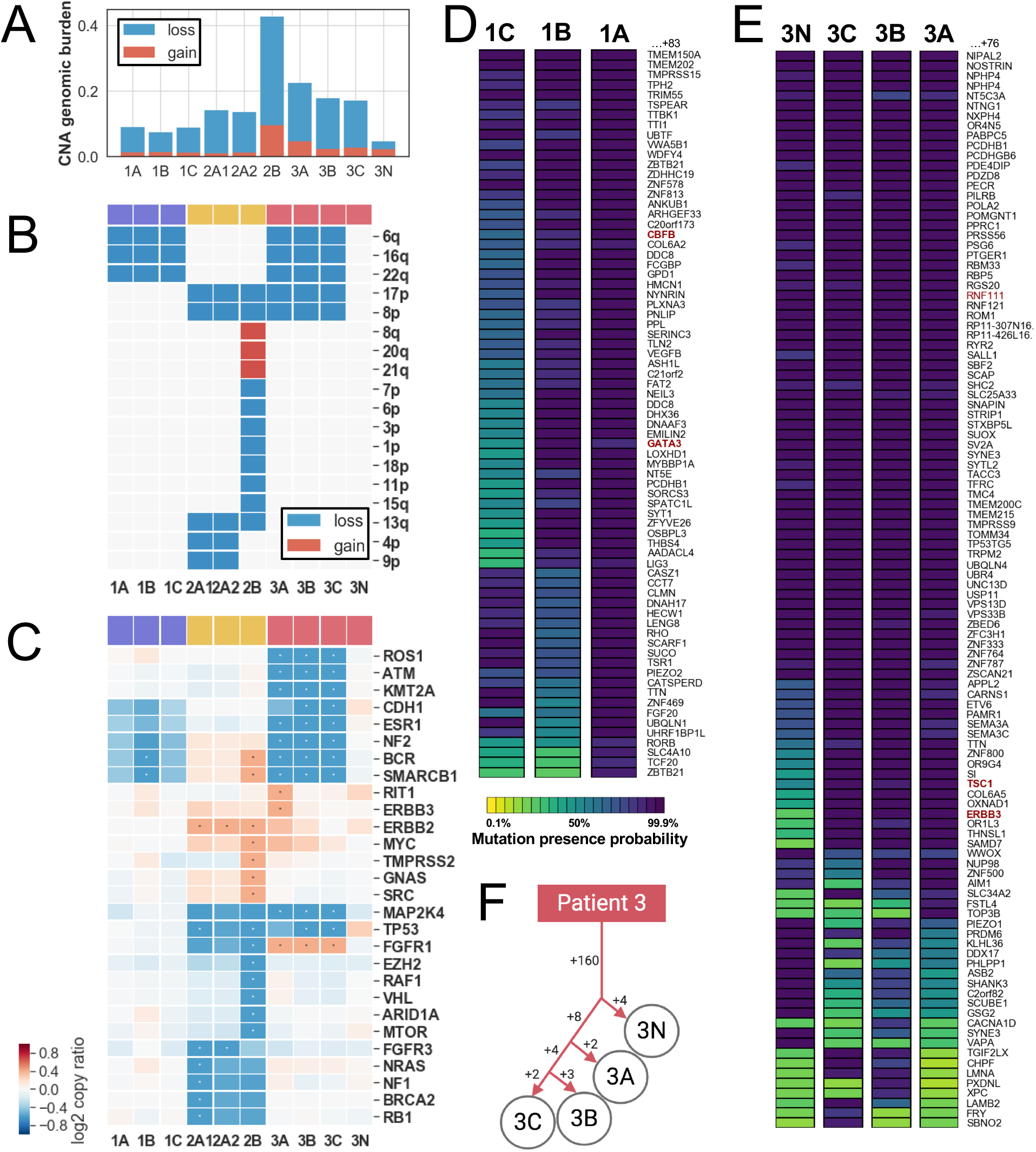
Mutational profile and clonal analysis from multi-region DNA sequencing of DCIS patients. **(A)** Fraction of genome involved in copy number losses (blue) or gains (red) for each sequenced region. **(B)** Chromosome arm copy number status in each sequenced region: lost (blue) or gained (red). (**C**) Cancer gene copy number (log_2_ ratio – blue red gradient). Genes from the cancer panel with copy number gain (log_2_(ratio)>0.4) or loss (log_2_(ratio)<-0.6) in at least one region, indicated with an asterisk, are displayed. **(D-E)** Bayesian probabilistic variant classification of selected high confidence somatic variants (represented by their cognate gene – rows) across all dissected regions of same patient (columns). Variants are shown for patient 1 (D) and patient 3 (E). The color gradient indicates the posterior probability of mutation presence in each region. Genes from the Cancer Gene Census are indicated in red font. **(F)** Maximum likelihood tree generated using somatic mutations identified in patient 3’s regions.

We next identified somatic mutations in each region of patient 1 and 3’s specimens. After additional quality filtering using cross-patient information, we identified between 18 and 154 somatic mutations per PML region, of which 14 to 108 were non-silent (Additional file 1: Table S7). The resulting mutational burden (0.46 – 3.97 mutations/Mb) is a range similar to what has been observed in invasive breast cancer [50]. The somatic mutations were then used to characterize the clonal relationships between regions. To account for uneven sequencing coverage between regions and the possibility of random allelic dropout, we used a maximum likelihood approach, comparing allelic fraction and total read depth of any mutated position in all regions of patients 1 and 3 (Figure 4D-E) [37]. Mutations in patient 1 were evaluated at 155 mutated positions, of which 141 were confidently identified in all 3 regions, and 14 were either missing or absent from one or more region (Additional file 1: Table S8). While a portion of these may be shared mutations may be germline variants, we identified a *GATA3* splice-site deletion in regions 1A and 1B previously observed in DCIS studies, disrupting a canonical splice site and for which the resulting transcript has been shown to lead to an abnormally high number of neoantigens [51,52]. Similarly, mutations in patient 3 were evaluated at 183 positions, 160 of which were shared by all 4 regions, including normal, and likely residual germline variants. The results were used to build a phylogenic tree illustrating the clonal relationship between regions (Figure 4F). Interestingly, region 3N is mutated at 4 positions not found in the PML regions. These could represent mutations acquired in aging normal tissue or residual germline variants lost in the PMLs [53–55]. The 3 PML regions gained an additional 8 mutations before diverging including an *ERBB3 Ile763Leu* substitution was exclusively observed in all 3 PML regions of patient 3. This mutation is predicted to be deleterious, possibly activating this uncommon driver of breast cancer [56,57] and may have contributed, in concert with *FGFR1* gain, to the clonal expansion observed in this patient. Region 3A gained an additional 2 mutations, while 3C and 3B shared an additional 4 and each gained between 1 and 3 mutations that were not shared. Overall, our analysis suggests that even in absence of normal DNA, and akin to experiments in large tumors, variants from multi-region sequencing can be used to trace evolutionary relationships between areas of pre-malignancy [37,58].

## Discussion

The results presented here demonstrate our ability to perform comprehensive mutational profiling from some of the most challenging clinical specimens with DNA in limited quantity – down to 3 ng – and of poor quality – highly degraded and chemically altered. In particular, this demonstration relied on a thorough benchmarking study using DNA from mirrored matching frozen vs FFPE specimens, which provided a real-world experimental framework to guide the process development.

We demonstrated that by utilizing a sequencing library preparation that uses a non-standard adapter ligation, we can drastically improve sequencing performance from these challenging specimens. A-tailing, together with transposon-mediated construction, is one of the most popular methods to prepare high-throughput sequencing libraries. While the latter necessitates longer DNA fragments and is not-suitable for highly degraded DNA, A-tailing has been broadly used in library preparation and is compatible with highly degraded specimens so long as DNA input is increased. We illustrated this limitation, analyzing targeted sequencing from limited dilutions and saw a drop in coverage and variant calling accuracy below 50 ng DNA input. Target coverage cannot be rescued by sequencing of a smaller panel, since the bottleneck resulting in lack of library complexity occurs prior to capture. For clinical reasons, to allow the thorough inspection of all tissue parts to formally exclude invasive disease, all pure PMLs are fixed in formalin and archived. Furthermore, areas of PML are typically small, limiting the quantity of material available for analysis. For the largest lesions, a labor-intensive dissection and pooling can increase the amount of DNA extracted but would preclude the study of their genetic heterogeneity. Similar to the benefit demonstrated on ancient DNA [41,43], we showed that an alternative library preparation using either single-strand and even more so for blunt-end ligation strategies considerably improved both coverage and consequently variant calling performances. Despite the recognized high efficiency of sticky-end ligation [59], end-repair and addition of an overhanging adenosine is likely the rate-limiting step on highly damaged DNA.

Another challenge we faced and addressed was calling mutations in absence of a matching normal DNA in both the test and PML DNA, likely leading to some ambiguous mutation classification [60]. The most useful PML specimens for biomarker studies are the ones with long follow-up, and aside from logistical challenges of collecting matching blood or saliva as part of routine clinical workflow, these traditional sources of normal DNA are generally not available for archival PML. The dissection of adjacent normal tissue, performed for one specimen in this study, can be sometimes used. While sufficient material can be found at the margin of the surgical specimen, the cellularity of the normal tissue will vary greatly between organs and histological context leading to even smaller quantities of DNA. In the breast, the normal ductal tree is poorly cellular in comparison to dysplastic and in situ proliferative lesions. In some instances, an area of high lymphocytic infiltration, for example at the location of a previous biopsy, can be used. Such histologically normal tissue are also formalin fixed and their mutational profiling presents similar, if not more, challenges than for PMLs. Furthermore, and as demonstrated elsewhere, some histologically normal specimens will contain a few somatic mutations at low-allelic fraction, resulting from early clonal selection, such as the few private mutations identified in sample 3N [53–55]. Nevertheless, the inclusion of such a matched normal DNA in the analysis and interpretation would greatly aid in the removal of residual germline DNA and additional sequencing artifacts. In absence of matching normal DNA, the parallel sequencing of a panel of unmatched normal DNA, from the same ethnic background and processed using the exact same protocol and analysis is recommended, especially in a clinical setting [60]. This was not available for this benchmarking study. Instead, we combined the use of a publicly obtained pool of normal with filtering for common variants using public databases following recommendation provided elsewhere [30,61]. This approach will miss rare germline variants, especially present in rare, or under-studied ethnicities. Previous studies have observed that tumor-only exomes may lead to ~300 residual germline variants after careful filtering [60,62]. Importantly, coding and deleterious germline variants could be a source of false positives. In our study, multi-region sampling provided additional information to help us classify mutations, and we determined that 155-160 mutations were shared and likely represented residual germline variants. This approach would however not remove the ambiguity for shared mutations and, in the event this mutation is an oncogenic driver common to all regions, additional validation steps may be required, including sequencing from a more distant region located on a separate tissue block, or comparison to its allelic fraction and copy number status in the bulk DNA.

Independent of the possible residual germline variants, we took specific steps to benchmark variant calling in highly degraded FFPE DNA, comparing the results to high quality variants called from the DNA of an adjacent frozen specimen. After carefully down-sampling the data to obtain the same number of raw sequencing reads – including *in silico* “replicates” from the deeper reference (frozen tissue) dataset itself – we identified two main mode of errors. First, a large number of false negative variants, leading to lower recall, were directly associated to the uneven and lower coverage depth, particularly in the low input FFPE AT library. The high and more even coverage observed in the BE libraries, for the same number of raw reads, remediated this issue, resulting in recall similar to the one expected from sampling bias observed in other studies [63]. Second, the false positives were mostly due to C to T substitutions as a consequence of formalin fixation, as previously observed. Such substitutions however remained at low allelic fraction and displayed strong strand bias, which could be remediated using stringent heuristic filtering. Alternate solutions have been described elsewhere to remove same or similar sequencing artifacts using machine learning [64] or relying on the precise substitution signature of the artifacts [65,66]. These would likely yield similar or superior results. DNA damage can also be repaired prior to library preparation altogether using cocktails of DNA repair enzymes, such as UDG and Fpg [67,68]. This strategy would decrease false positive mutations [69] but typically require more than 5 ng FFPE DNA and the extra enzymatic step would likely add bottleneck and decrease library complexity.

Of the 3 DCIS patients studied, all displayed chromosomal copy number changes previously observed in DCIS, leading to losses of known tumor suppressors (*TP53*) or gains of known oncogenes *(ERBB2* or *FGFR1).* In 2/3 patients, we observed a comparable number of somatic mutations to pure DCIS previously studies [70,71]. The identification of only two known or likely breast cancer driver mutations *(GATA3* and *ERBB3)* in two of the specimen was not surprising, as these specimens represent pure DCIS lacking any invasive component, and thus should bear less genomic resemblance to breast cancer [70]. Interestingly *GATA3* has been reported as mutated at a higher frequency in pure DCIS than in invasive cancer, suggesting a negative selection during transition to invasive cancer [52]. This particular splice-site mutation produces an alternative transcript resulting in a high numbers of neoantigens, perhaps subjecting mutated lesions to a more effective immune-surveillance [51]. The relative contribution of gene copy number alterations *vs* somatic mutations to cellular proliferation and clonal selection in normal and pre-malignant tissue is an active field of study [53–55] and progress in this field will require multi-modal molecular profiling approaches compatible with small amounts of archived tissue, such as the one described here.

Beyond the identification of drivers of growth and proliferation, the proposed approach can help measure the genetic heterogeneity in PML lesions. In breast DCIS, phenotypic heterogeneity associated with subtype and grade can co-exist within a specimen [72]. Additionally, immunostaining of key markers has revealed spatial heterogeneity within a duct and between ducts of a patient [73]. But the mutational landscape underlying such heterogeneity has not been thoroughly studied in pure DCIS at a genome-wide level for the technical reasons mentioned herein. Multi-region assessment of karyotypes, select gene copy number [74] and mitochondrial mutations [75] suggested significant heterogeneity in DCIS but its clinical significance and association with other progression risk factors has not been assessed. Copynumber heterogeneity has also been observed via single-cell sequencing from frozen DCIS patient specimens, albeit in the presence of an invasive lesion [15]. A similar approach has been developed in archived tissue specimens, but none have been used in large studies or in pure DCIS [76]. While single-cell sequencing may be able to scale up – both number of cells and number of samples, it cannot yet identify point mutations with high sensitivity. Hence, in the context of a clinical study, multi-region sequencing enabled by LCM may be preferable as it would increase the accuracy of the clonal evolution and enable the identification of driver mutations and mutational signatures [77]. Moreover, the results of multi-region genomic profiling would enable us to place somatic mutations and copy number alterations in the context of the surrounding extra-cellular matrix or stromal composition obtained via imaging and morphological studies, providing a granular view of the premalignant landscape as aspired by the pre-cancer atlas [78].

## Conclusions

We present both methodological and analytical advances enabling targeted sequencing from small, archived PML lesions. Specifically, we showed that blunt-end ligation of the sequencing adapter to the fragmented and damaged DNA was key to increasing the quality of the exome sequencing from a minute amount of DNA extracted from archived specimens. A subsequent multi-caller analysis strategy, complemented by dedicated filters, identified somatic mutations, eliminating false-positive and residual germline mutations – two critical hurdles in the mutational profiling of PML. Importantly, the approach was developed on real world samples using a well-characterized test sample and further validated on micro-dissected PML regions. These advances now enable mutational profiling of PML lesions, unlocking the untapped registry of archived PML specimens to comprehensively investigate their molecular heterogeneity and assess the contribution of somatic DNA alterations to cellular proliferation and progression to invasive cancer. These tools may provide clues as to which PML should be treated to prevent invasive cancer and which can be left alone.

## List of abbreviations

BE: Blunt-end
SS: Single-strand
AT: A-tailing
DCIS: Ductal carcinoma in situ
LCM: Laser capture microdissection
IDC: Invasive ductal carcinoma
FFPE: Formalin-fixed paraffin embedded
LOH: Loss of heterozygosity
VAF: Variant allelic frequency
PML: Pre-malignant lesion
CNA: Copy number alteration
Cov20: Fraction of base pairs covered at 20x or more

## Declarations

### Ethics approval and consent to participate

The retrospective use of clinical specimen was approved by the UC San Francisco Institutional Review Board and UC San Diego Institutional Review Board who granted a waiver of informed consent.

### Consent for publication

Not applicable

### Availability of data and materials

Configuration of bcbio pipeline is available publicly on GitHub: https://github.com/danielanach/bcbio-ffpe-no-normal-config/blob/master/bcbio_ffpe-no-normal.yaml. The raw sequencing data for the test specimen have been deposited with links to BioProject accession number PRJNA615755 in the NCBI BioProject database. The raw sequencing data from the DCIS patients has been deposited to the NCBI dbGaP (release pending).

### Authors’ contributions

SB and FH selected specimen and performed pathology annotation, EZ and TOK identified breast cancer patients and reviewed associated clinical data. JS prepared the DNA samples, KJ and HY performed the targeted sequencing. KJ, DN, OH designed the test study, OH, GH, LE, SB designed the validation study. AO performed the RNA-seq analysis. DN and OH performed the analysis and wrote the manuscript. All authors approved the final manuscript.

## Acknowledgements

We express our gratitude to members of the Moores Cancer Center Biotechnology and Tissue Technology Shared Resource: Drs Molinolo, Kaushal, Estrada and Kimberly McIntyre for their technical assistance. All sequencing was conducted at the IGM Genomics Center, University of California, San Diego, La Jolla, CA.

## References

1. Esserman LJ, Thompson IM, Reid B. Overdiagnosis and overtreatment in cancer: An opportunity for improvement. JAMA - J Am Med Assoc. 2013;310:797–8.

2. Siegel RL, Miller KD, Jemal A. Cancer statistics, 2020. CA Cancer J Clin. 2020;70:7–30.

3. Marmot M, Altman DG, Cameron DA, Dewar JA, Thompson SG, Wilcox M. The benefits and harms of breast cancer screening: An independent review. Lancet. 2012;380:1778–86.

4. Bleyer A, Welch HG. Effect of three decades of screening mammography on breast-cancer incidence. N Engl J Med. 2012;367:1998–2005.

5. Menon R, Deng M, Boehm D, Braun M, Fend F, Boehm D, et al. Exome enrichment and SOLiD sequencing of formalin fixed paraffin embedded (FFPE) prostate cancer tissue. Int J Mol Sci. 2012;13:8933–42.

6. Hedegaard J, Thorsen K, Lund MK, Hein AMK, Hamilton-Dutoit SJ, Vang S, et al. Nextgeneration sequencing of RNA and DNA isolated from paired fresh-frozen and formalin-fixed paraffin-embedded samples of human cancer and normal tissue. PLoS One. 2014;9:e98187.

7. Allen EM Van, Wagle N, Stojanov P, Perrin DL, Cibulskis K, Marlow S, et al. Whole-exome sequencing and clinical interpretation of FFPE tumor samples to guide precision cancer medicine. Nat Med. 2014;20:682–8.

8. Munchel S, Hoang Y, Zhao Y, Cottrell J, Klotzle B, Godwin AK, et al. Targeted or whole genome sequencing of formalin fixed tissue samples: Potential applications in cancer genomics. Oncotarget. 2015;6:25943–61.

9. Astolfi A, Urbini M, Indio V, Nannini M, Genovese CG, Santini D, et al. Whole exome sequencing (WES) on formalin-fixed, paraffin-embedded (FFPE) tumor tissue in gastrointestinal stromal tumors (GIST). BMC Genomics. 2015;16:892.

10. Miron A, Varadi M, Carrasco D, Li H, Luongo L, Kim HJ, et al. PIK3CA mutations in in situ and invasive breast carcinomas. Cancer Res. 2010;70:5674–8.

11. Sontag L, Axelrod DE. Evaluation of pathways for progression of heterogeneous breast tumors. J Theor Biol. 2005;232:179–89.

12. Yates LR, Gerstung M, Knappskog S, Desmedt C, Gundem G, Van Loo P, et al. Subclonal diversification of primary breast cancer revealed by multiregion sequencing. Nat Med. 2015;21:751–9.

13. Newburger DE, Kashef-Haghighi D, Weng Z, Salari R, Sweeney RT, Brunner AL, et al. Genome evolution during progression to breast cancer. Genome Res. 2013;23:1097–108.

14. Oikawa M, Yano H, Matsumoto M, Otsubo R, Shibata K, Hayashi T, et al. A novel diagnostic method targeting genomic instability in intracystic tumors of the breast. Breast Cancer. 2015;22:529–35.

15. Casasent AK, Schalck A, Gao R, Sei E, Long A, Pangburn W, et al. Multiclonal Invasion in Breast Tumors Identified by Topographic Single Cell Sequencing. Cell. 2018;72:205–17.

16. Arreaza G, Qiu P, Pang L, Albright A, Hong LZ, Marton MJ, et al. Pre-analytical considerations for successful next-generation sequencing (NGS): Challenges and opportunities for formalin-fixed and paraffin-embedded tumor tissue (FFPE) samples [Internet]. Int. J. Mol. Sci. 2016. Available from: http://dx.doi.org/10.3390/ijms17091579

17. Do H, Dobrovic A. Sequence artifacts in DNA from formalin-fixed tissues: Causes and strategies for minimization. Clin Chem. 2015;61:64–71.

18. Foley JW, Zhu C, Jolivet P, Zhu SX, Lu P, Meaney MJ, et al. Gene expression profiling of single cells from archival tissue with laser-capture microdissection and Smart-3SEQ. Genome Res. 2019;

19. Illumina. bcl2fastq Conversion Software v1.8.4. Illumina. 2018.

20. Guimera RV. bcbio-nextgen: Automated, distributed next-gen sequencing pipeline. EMBnet.journal [Internet]. 2012;17:30. Available from: http://journal.embnet.org/index.php/embnetjournal/article/view/286

21. Li H. Seqtk. GitHub. 2015.

22. Didion JP, Martin M, Collins FS. Atropos: Specific, sensitive, and speedy trimming of sequencing reads. PeerJ. 2017;5:e3720.

23. Li H, Durbin R. Fast and accurate short read alignment with Burrows-Wheeler transform. Bioinformatics. 2009;25:1754–60.

24. Broad Institute. Picard tools [Internet]. https://broadinstitute.github.io/picard/. 2016. Available from: https://broadinstitute.github.io/picard/ http://broadinstitute.github.io/picard/

25. Talevich E, Shain AH, Botton T, Bastian BC. CNVkit: Genome-Wide Copy Number Detection and Visualization from Targeted DNA Sequencing. PLoS Comput Biol. 2016;2:e1004873.

26. Olshen AB, Bengtsson H, Neuvial P, Spellman PT, Olshen RA, Seshan VE. Parent-specific copy number in paired tumor-normal studies using circular binary segmentation. Bioinformatics. 2011;27:2038–46.

27. Lai Z, Markovets A, Ahdesmaki M, Chapman B, Hofmann O, Mcewen R, et al. VarDict: A novel and versatile variant caller for next-generation sequencing in cancer research. Nucleic Acids Res. 2016;44:e108.

28. Benjamin D, Sato T, Cibulskis K, Getz G, Stewart C, Lichtenstein L. Calling Somatic SNVs and Indels with Mutect2. bioRxiv. 2019;

29. Cingolani P, Platts A, Wang LL, Coon M, Nguyen T, Wang L, et al. A program for annotating and predicting the effects of single nucleotide polymorphisms, SnpEff. Fly (Austin) [Internet]. 2012;6:80–92. Available from: http://www.tandfonline.com/doi/abs/10.4161/fly.19695

30. bcbio tumor-only. Available from: https://bcbio.wordpress.com/2015/03/05/cancerval/

31. 1000 Genomes Project Consortium. An integrated map of genetic variation. Nature. 2012;491:56–65.

32. Exome Aggregate Consortium. ExAC Browser. Online. 2016.

33. Karczewski KJ, Francioli LC, Tiao G, Cummings BB, Alföldi J, Wang Q, et al. Variation across 141,456 human exomes and genomes reveals the spectrum of loss-of-function intolerance across human protein-coding genes. bioRxiv. 2019;

34. Forbes SA, Beare D, Boutselakis H, Bamford S, Bindal N, Tate J, et al. COSMIC: Somatic cancer genetics at high-resolution. Nucleic Acids Res. 2017;45:D777–83.

35. Landrum MJ, Lee JM, Benson M, Brown GR, Chao C, Chitipiralla S, et al. ClinVar: Improving access to variant interpretations and supporting evidence. Nucleic Acids Res. 2018;46:D1062–7.

36. Cleary JG, Braithwaite R, Gaastra K, Hilbush BS, Inglis S, Irvine SA, et al. Joint variant and de novo mutation identification on pedigrees from high-throughput sequencing data. J Comput Biol. 2014;21:405–19.

37. Reiter JG, Makohon-Moore AP, Gerold JM, Bozic I, Chatterjee K, Iacobuzio-Donahue CA, et al. Reconstructing metastatic seeding patterns of human cancers. Nat Commun. 2017;8:14114.

38. Levy E, Marty R, Gárate Calderón V, Woo B, Dow M, Armisen R, et al. Immune DNA signature of T-cell infiltration in breast tumor exomes. Sci Rep. 2016;6:30064.

39. Troll CJ, Kapp J, Rao V, Harkins KM, Cole C, Naughton C, et al. A ligation-based singlestranded library preparation method to analyze cell-free DNA and synthetic oligos. BMC Genomics. 2019;20:1023.

40. Bennett CW, Berchem G, Kim YJ, El-Khoury V. Cell-free DNA and next-generation sequencing in the service of personalized medicine for lung cancer. Oncotarget. 2016;7:71013–35.

41. Bennett EA, Massilani D, Lizzo G, Daligault J, Geigl EM, Grange T. Library construction for ancient genomics: Single strand or double strand? Biotechniques. 2014;

42. Dabney J, Knapp M, Glocke I, Gansauge MT, Weihmann A, Nickel B, et al. Complete mitochondrial genome sequence of a Middle Pleistocene cave bear reconstructed from ultrashort DNA fragments. Proc Natl Acad Sci U S A. 2013;110:15758–63.

43. Gansauge MT, Meyer M. Single-stranded DNA library preparation for the sequencing of ancient or damaged DNA. Nat Protoc. 2013;8:737–48.

44. Sundaram AYM, Hughes T, Biondi S, Bolduc N, Bowman SK, Camilli A, et al. A comparative study of ChIP-seq sequencing library preparation methods. BMC Genomics. 2016;17:816.

45. Ma S, Hsieh YP, Ma J, Lu C. Low-input and multiplexed microfluidic assay reveals epigenomic variation across cerebellum and prefrontal cortex. Sci Adv. 2018;4:eaar8187.

46. Costello M, Pugh TJ, Fennell TJ, Stewart C, Lichtenstein L, Meldrim JC, et al. Discovery and characterization of artifactual mutations in deep coverage targeted capture sequencing data due to oxidative DNA damage during sample preparation. Nucleic Acids Res. 2013;41:e67.

47. Kader T, Goode DL, Wong SQ, Connaughton J, Rowley SM, Devereux L, et al. Copy number analysis by low coverage whole genome sequencing using ultra low-input DNA from formalin-fixed paraffin embedded tumor tissue. Genome Med. 2016;8:121.

48. Rieber N, Bohnert R, Ziehm U, Jansen G. Reliability of algorithmic somatic copy number alteration detection from targeted capture data. Bioinformatics. 2017;33:2791–2798.

49. Reis-Filho JS, Lakhani SR. The diagnosis and management of pre-invasive breast disease Genetic alterations in pre-invasive lesions. Breast Cancer Res. 2003;313-9:313–9.

50. Weinstein JN, Collisson EA, Mills GB, Shaw KRM, Ozenberger BA, Ellrott K, et al. The cancer genome atlas pan-cancer analysis project. Nat Genet. 2013;45:1113–20.

51. Jayasinghe RG, Cao S, Gao Q, Wendl MC, Vo NS, Reynolds SM, et al. Systematic Analysis of Splice-Site-Creating Mutations in Cancer. Cell Rep. 2018;

52. Pang JMB, Savas P, Fellowes AP, Mir Arnau G, Kader T, Vedururu R, et al. Breast ductal carcinoma in situ carry mutational driver events representative of invasive breast cancer. Mod Pathol. 2017;

53. Martincorena I, Fowler JC, Wabik A, Lawson ARJ, Abascal F, Hall MWJ, et al. Somatic mutant clones colonize the human esophagus with age. Science (80-). 2018;

54. Martincorena I, Roshan A, Gerstung M, Ellis P, Van Loo P, McLaren S, et al. High burden and pervasive positive selection of somatic mutations in normal human skin. Science (80-). 2015;348:880–6.

55. Salk JJ, Loubet-Senear K, Maritschnegg E, Valentine CC, Williams LN, Higgins JE, et al. Ultra-Sensitive TP53 Sequencing for Cancer Detection Reveals Progressive Clonal Selection in Normal Tissue over a Century of Human Lifespan. Cell Rep. 2019;28:132–44.

56. Bose R, Kavuri SM, Searleman AC, Shen W, Shen D, Koboldt DC, et al. Activating HER2 mutations in HER2 gene amplification negative breast cancer. Cancer Discov. 2013;3:224–37.

57. Mujoo K, Choi BK, Huang Z, Zhang N, An Z. Regulation of ERBB3/HER3 signaling in cancer. Oncotarget. 2014;

58. Gerlinger M, Horswell S, Larkin J, Rowan AJ, Salm MP, Varela I, et al. Genomic architecture and evolution of clear cell renal cell carcinomas defined by multiregion sequencing. Nat Genet. 2014;

59. Quail MA, Kozarewa I, Smith F, Scally A, Stephens PJ, Durbin R, et al. A large genome center’s improvements to the Illumina sequencing system. Nat Methods. 2008;

60. Jones S, Anagnostou V, Lytle K, Parpart-Li S, Nesselbush M, Riley DR, et al. Personalized genomic analyses for cancer mutation discovery and interpretation. Sci Transl Med. 2015;362:911.

61. Kalatskaya I, Trinh QM, Spears M, McPherson JD, Bartlett JMS, Stein L. ISOWN: Accurate somatic mutation identification in the absence of normal tissue controls. Genome Med. 2017;9:59.

62. Garofalo A, Sholl L, Reardon B, Taylor-Weiner A, Amin-Mansour A, Miao D, et al. The impact of tumor profiling approaches and genomic data strategies for cancer precision medicine. Genome Med. 2016;9:51.

63. Chen Z, Yuan Y, Chen X, Chen J, Lin S, Li X, et al. Systematic comparison of somatic variant calling performance among different sequencing depth and mutation frequency. Sci Rep. 2020;10:3501.

64. Anzar I, Sverchkova A, Stratford R, Clancy T. NeoMutate: An ensemble machine learning framework for the prediction of somatic mutations in cancer. BMC Med Genomics. 2019;12:63.

65. Yost SE, Smith EN, Schwab RB, Bao L, Jung H, Wang X, et al. Identification of high-confidence somatic mutations in whole genome sequence of formalin-fixed breast cancer specimens. Nucleic Acids Res. 2012;40:e107.

66. Kim H, Lee AJ, Lee J, Chun H, Ju YS, Hong D. FIREVAT: Finding reliable variants without artifacts in human cancer samples using etiologically relevant mutational signatures. Genome Med. 2019;11:81.

67. Hofreiter M. DNA sequences from multiple amplifications reveal artifacts induced by cytosine deamination in ancient DNA. Nucleic Acids Res. 2001;29:4793–9.

68. Tchou J, Grollman AP. Repair of DNA containing the oxidatively-damaged base, 8-oxoguanine. Mutat Res Toxicol. 1993;299:277–87.

69. Chen L, Liu P, Evans TC, Ettwiller LM. DNA damage is a pervasive cause of sequencing errors, directly confounding variant identification. Science (80-). 2017;355:752–6.

70. Kim SY, Jung S-H, Kim MS, Baek I-P, Lee SH, Kim T-M, et al. Genomic differences between pure ductal carcinoma in situ and synchronous ductal carcinoma in situ with invasive breast cancer. Oncotarget. 2015;6:7597–607.

71. Pareja F, Brown DN, Lee JY, Da Cruz Paula A, Selenica P, Bi R, et al. Whole-Exome Sequencing Analysis of the Progression from Non-Low Grade Ductal Carcinoma &lt;em&gt;In Situ&lt;/em&gt; to Invasive Ductal Carcinoma. Clin Cancer Res [Internet]. 2020;clincanres.2563.2019. Available from: http://clincancerres.aacrjournals.org/content/early/2020/03/27/1078-0432.CCR-19-2563.abstract

72. Sinha VC, Piwnica-Worms H. Intratumoral Heterogeneity in Ductal Carcinoma In Situ: Chaos and Consequence. J Mammary Gland Biol Neoplasia. 2018;23:191–205.

73. Gerdes MJ, Gökmen-Polar Y, Sui Y, Pang AS, Laplante N, Harris AL, et al. Single-cell heterogeneity in ductal carcinoma in situ of breast. Mod Pathol. 2018;31:406–17.

74. Chin K, De Solorzano CO, Knowles D, Jones A, Chou W, Rodriguez EG, et al. In situ analyses of genome instability in breast cancer. Nat Genet. 2004;36:984–8.

75. Foschini MP, Morandi L, Leonardi E, Flamminio F, Ishikawa Y, Masetti R, et al. Genetic clonal mapping of in situ and invasive ductal carcinoma indicates the field cancerization phenomenon in the breast. Hum Pathol. 2013;44:1310–9.

76. Martelotto LG, Baslan T, Kendall J, Geyer FC, Burke KA, Spraggon L, et al. Whole-genome single-cell copy number profiling from formalin-fixed paraffin-embedded samples. Nat Med. 2017;23:376–85.

77. Alexandrov LB, Stratton MR. Mutational signatures: The patterns of somatic mutations hidden in cancer genomes. Curr Opin Genet Dev. 2014;24:52–60.

78. Srivastava S, Ghosh S, Kagan J, Mazurchuk R, Boja E, Chuaqui R, et al. The Making of a PreCancer Atlas: Promises, Challenges, and Opportunities. Trends in Cancer. 2019;4:523–36.

