## Additional File 2 for "Mutational profiling of micro-dissected pre-malignant lesions from archived specimens"

# S1

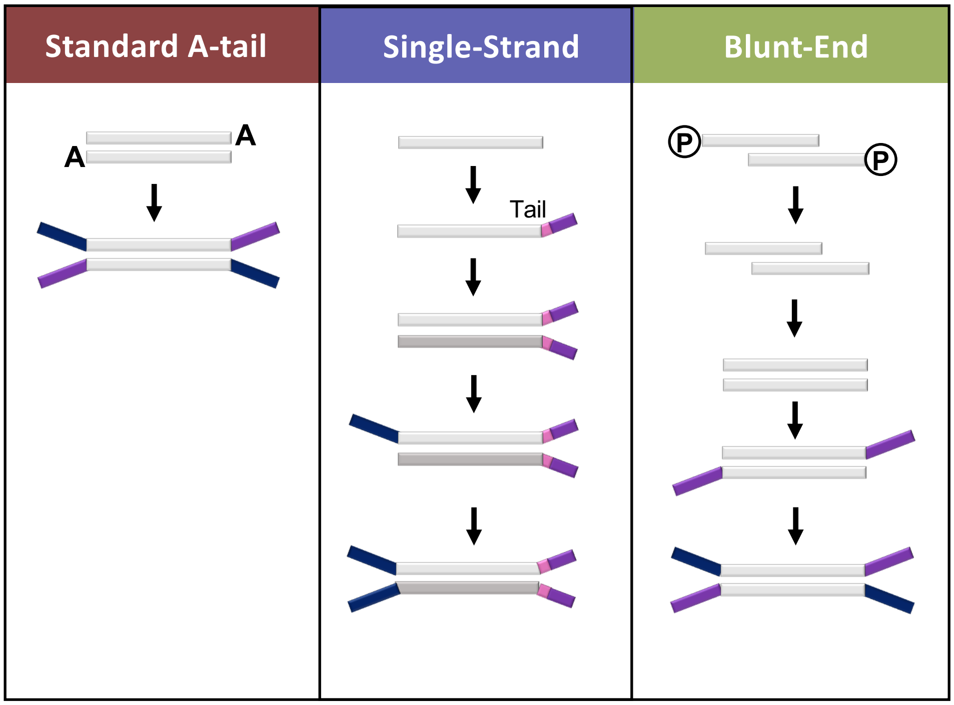


**Figure S1. Schematic overview of adapter ligation across library preparation strategies.** Purple and blue colors correspond to sequences containing Illumina P5 and P7 adapter sequences respectively.

# S2

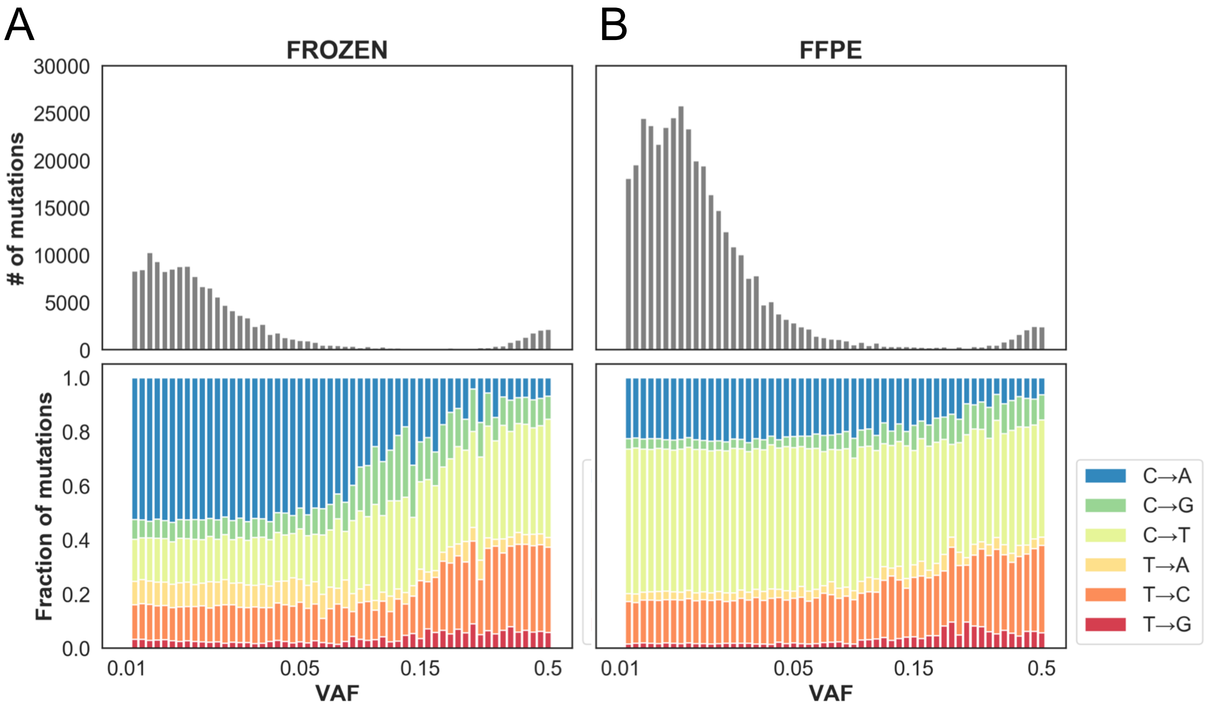


**Figure S2. Raw nucleotide substitution evaluation. (A-B)** The number (top) and substitution pattern (bottom) at variable variant allelic fractions (VAF - x-axis) observed in test specimen (200 ng DNA input) sequenced with AT library strategy across whole exome for frozen sample (A) and mirrored FFPE (B). Raw SNV substitutions were identified from VarDict output in absence of any filters. The proportion of each substitution is shown for each VAF bin in the bottom panels.

# S3

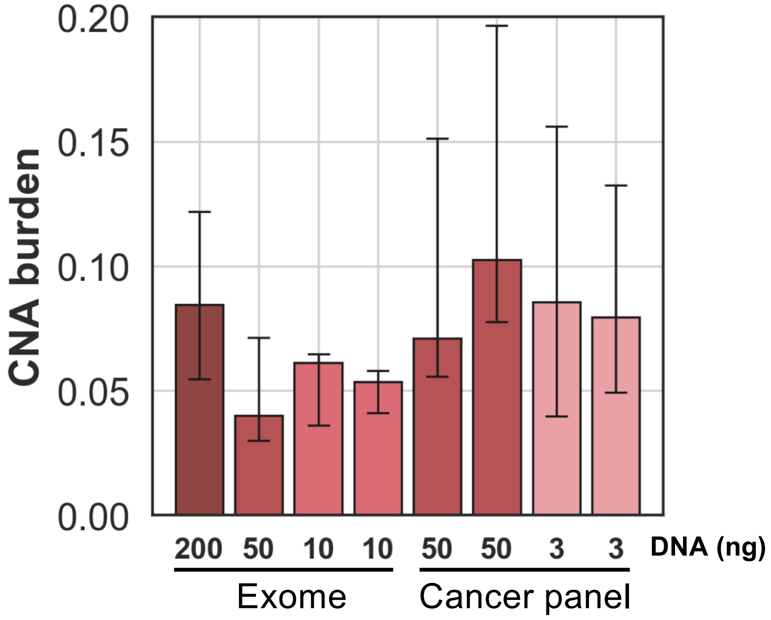


**Figure S3. Copy number burden estimates for AT libraries with varying DNA input.** Copy number alteration (CNA) burden was approximated as the fraction of genome in a CNA (-$\log_2$ copy<-0.3 or >0.3). Error bars represent CNA burden computed with either the upper or lower bounds of the 95% confidence interval around the $\log_2$ copy ratio of each segment .

# S4

A

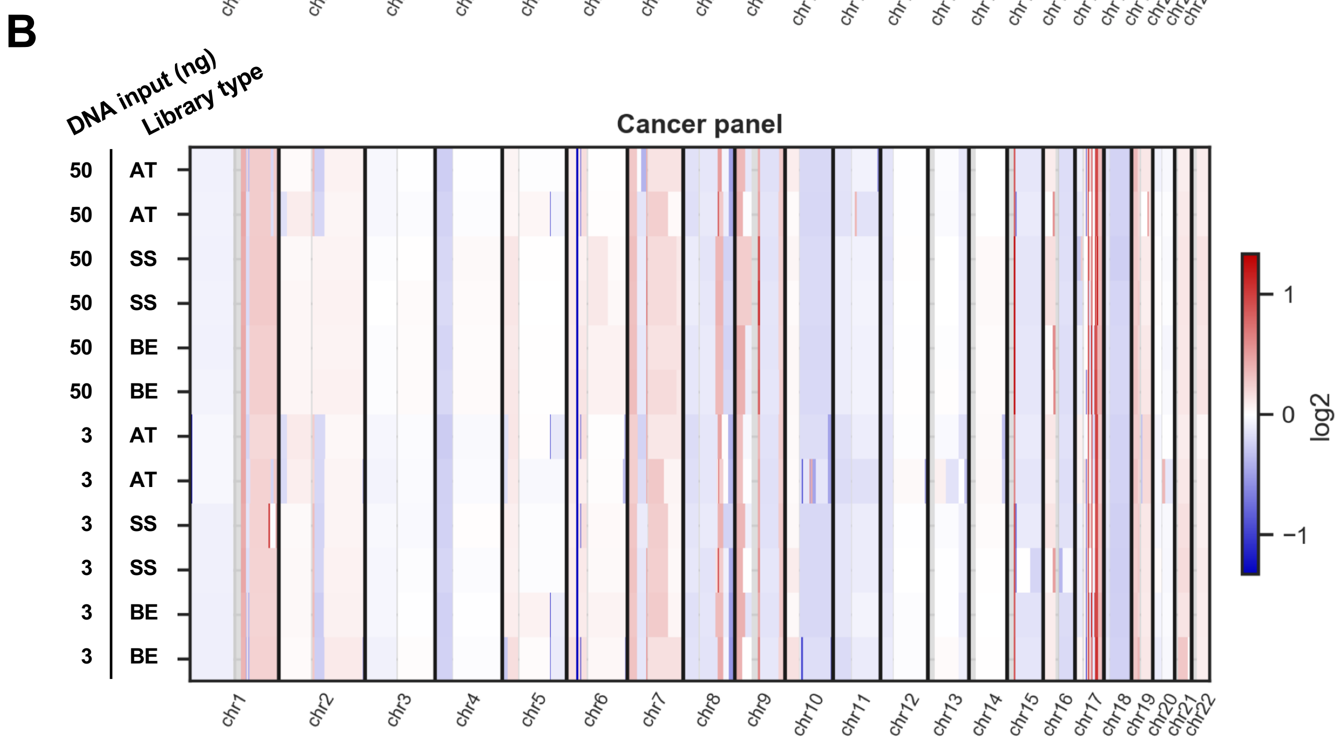


**Figure S4. Genome wide copy number profile across library preparation strategy and DNA input amount.** $Log_2$ copy ratio across entire genome for samples prepared with exome (A) and cancer panel (B).

# S5

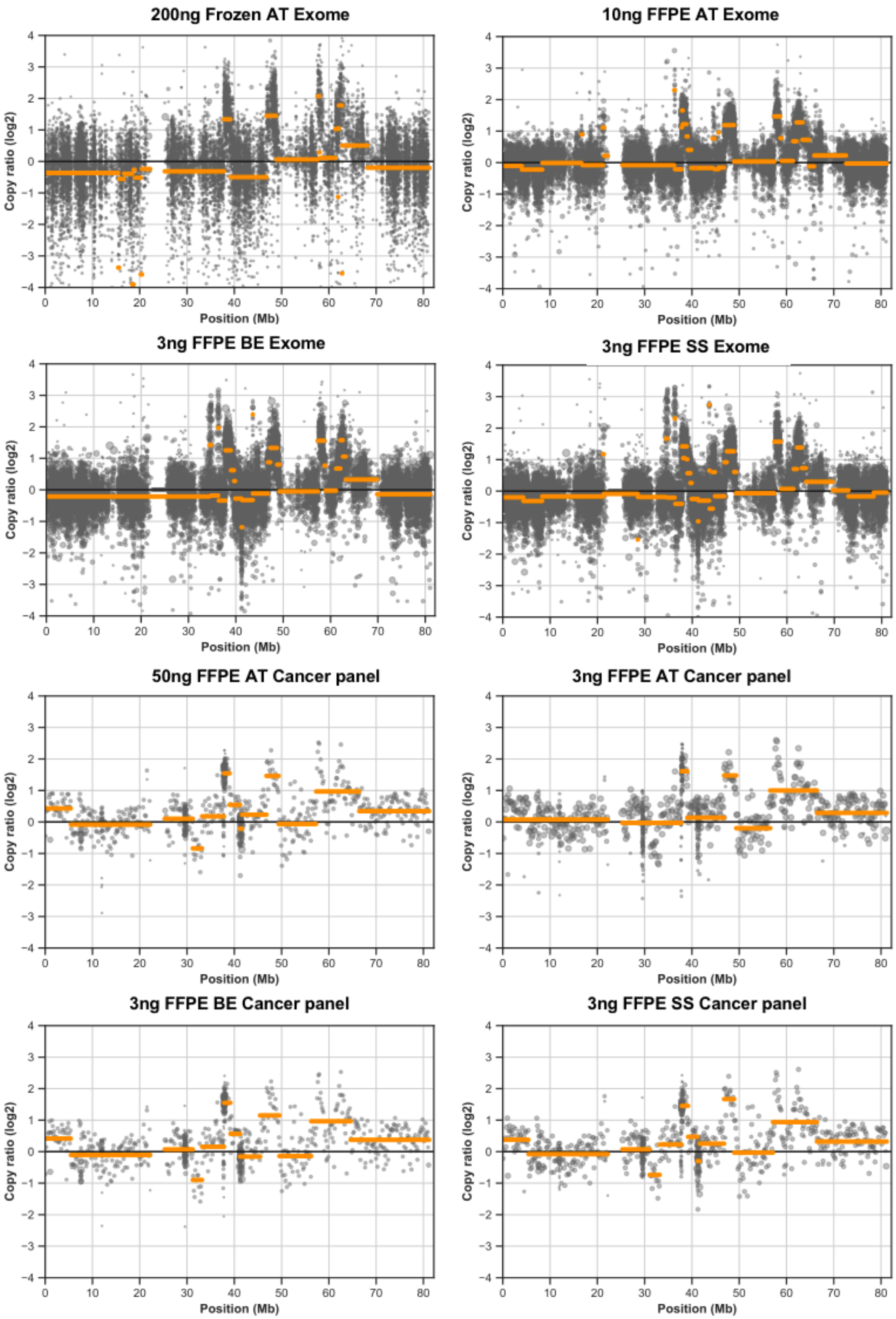


**Figure S5. Copy number profile of chromosome 17 across library preparation strategies and DNA input amount in FFPE test specimen.** Scatter plots show log2 copy ratios for bins (grey) and segments (orange). Library preparation strategy and DNA input amount are indicated above each panel

# S6

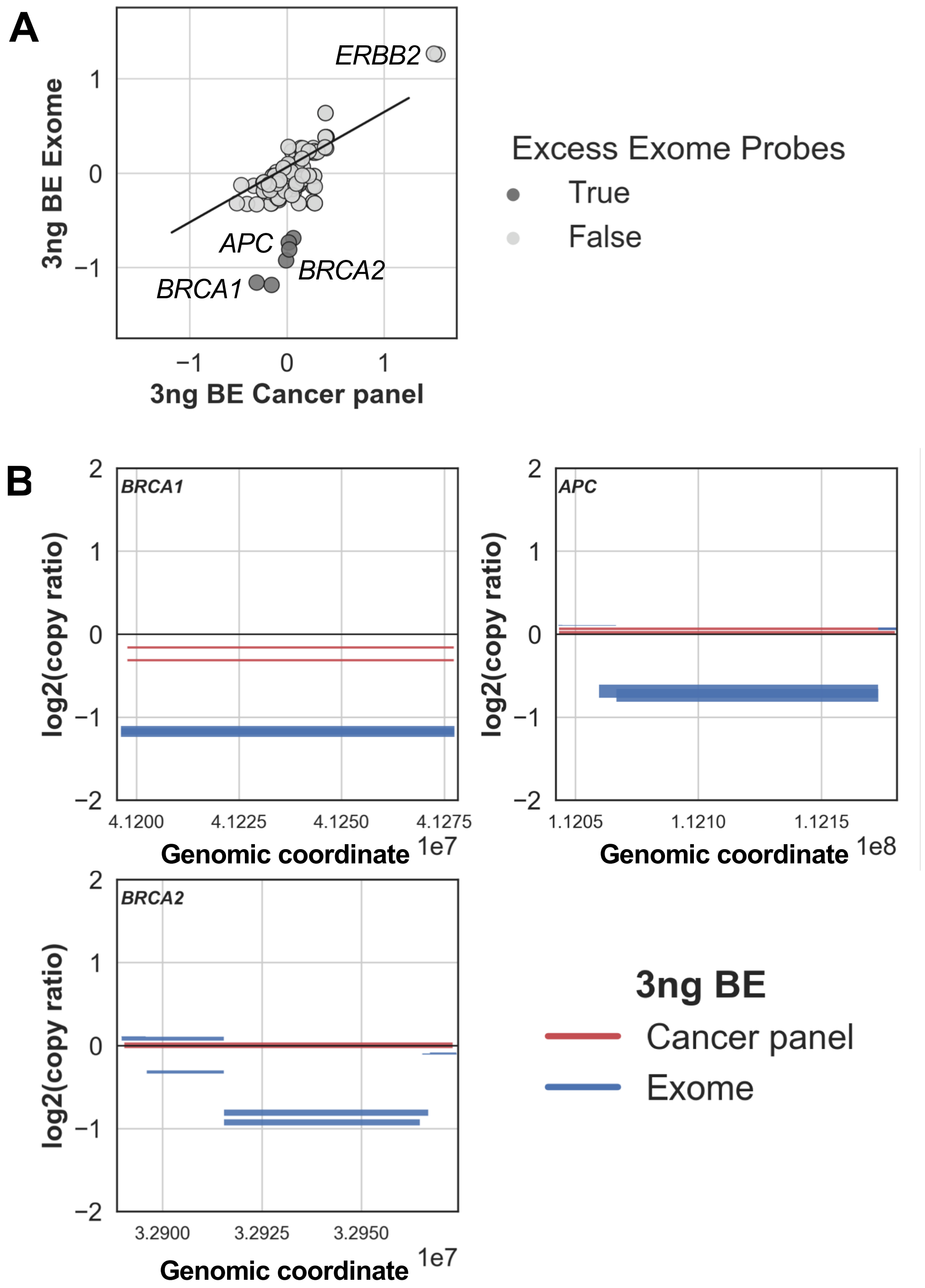


**Figure S6. Discordant copy number ratio between exome and cancer panel in low input BE library preparation strategy. (A)** Comparison of log2 copy number ratio observed in cancer panel (x-axis) and exome (y-axis) for 98 cancer genes. Genes covered by excess number of probes (>6x) in exome as compared to the cancer panel are colored in dark gray. **(B)** Copy number levels (log2 ratio - y axis) of genomic segments overlapping genes with discordant copy numbers (*BRCA1, BRCA2, APC*) between exome (blue) and cancer panel (red). All experimental replicates are displayed. Line thickness indicates the confidence level (thick=high) of the segment called.

# S7

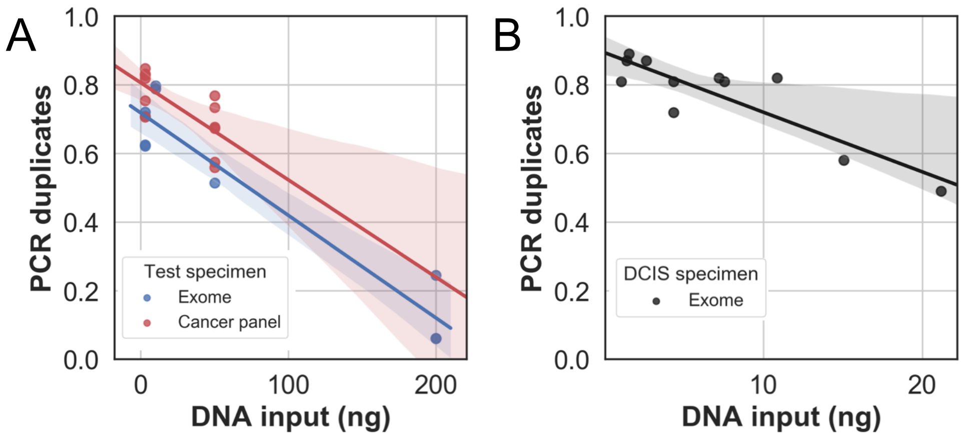


**Figure S7. PCR duplicate rate as a function of DNA input amount.** Fraction of PCR duplicate reads on y-axis as a function of DNA input on x-axis ($p$ = 1.4e-05) in test FFPE specimen (**A**) and for DCIS specimen ($p$ = 3e-03) (**B**).

# S8

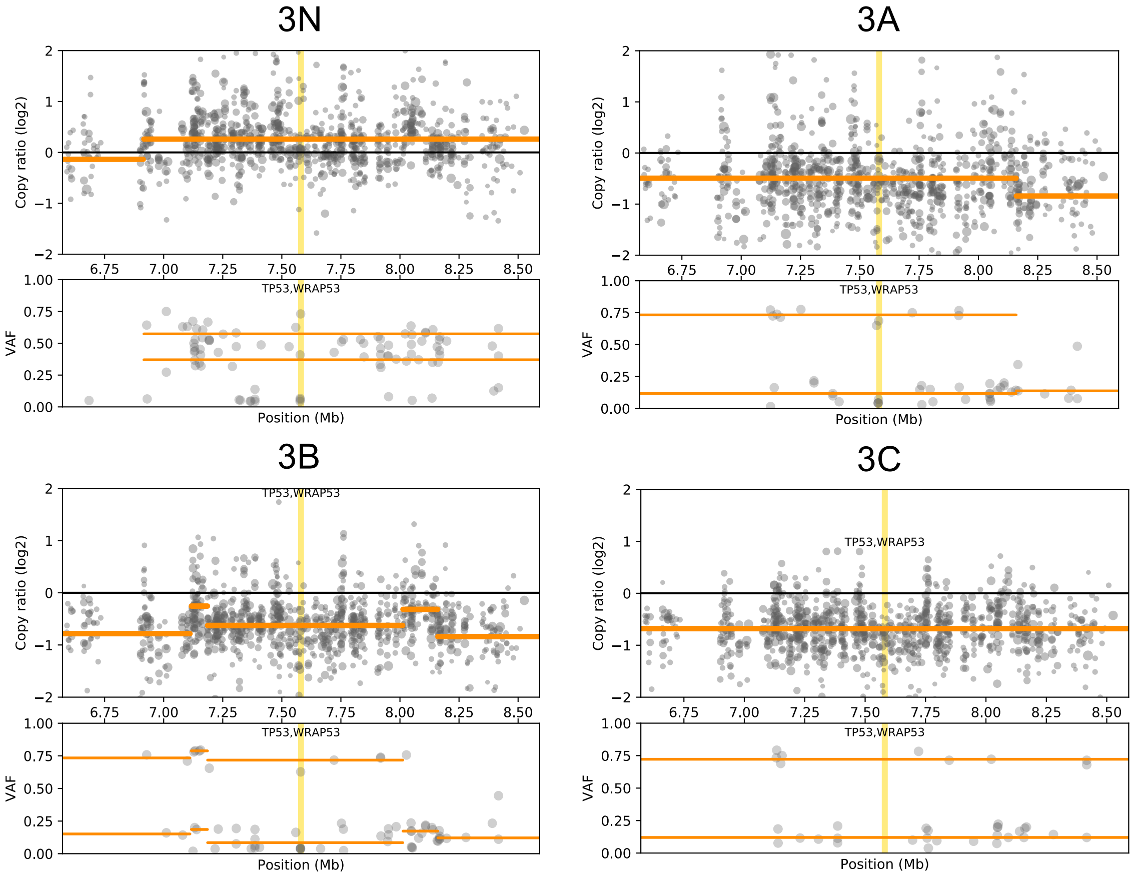


**Figure S8. *TP53* LOH in regions of patient 3.** Scatter plot of log2 copy number ratio (upper panels) and B-allele frequency (lower panels) of coverage bins (grey dots) and resulting genomic segments (orange lines) in a window of 2 Mb around TP53 (yellow stripe).

# S9

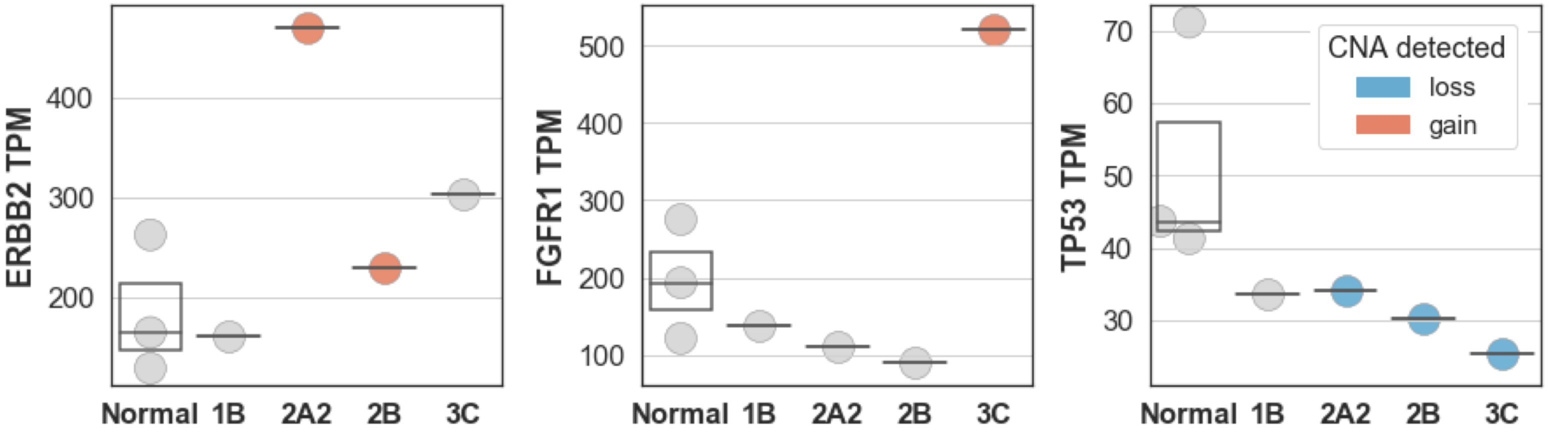


**Figure S9. Expression level of selected genes affected with CNA.** Normalized read counts, transcripts per million (TPM), from SMART-3Seq expression profiling of select DCIS regions for *ERBB2* (left), *FGFR1* (middle) and *TP53* (right), and as compared to unrelated normal dissected breast epithelium. Specimen in which a gene with a copy number gain was detected are shown in red, copy number loss in blue and, copy neutral in gray.
